# A Computational Study for Identifying Agronomically Essential Biomarkers Using nsSNP Data of Korean Soybeans

**DOI:** 10.1101/2020.06.20.161208

**Authors:** Yongwoon Jung, Nahyun You, Sangyong Nam

## Abstract

Soybean is a highly nutritious legume grown globally as a food and feed crop. An examination of a collection of 10 cultivated and 6 wild Korean soybean varieties showed that there is phenotypic variability notable in different soybeans. Therefore, to develop a list of biomarker candidates useful for growing soybeans of better quality and quantity, The genes of 16 Korean soybean varieties were compared with those of the reference Glycine max var. Williams 82. The comparison was made through gene sequencing to facilitate selection of nsSNPs. The objective of the study was to find out the structural and functional variations caused by nsSNPs and discuss whether the collection of Korean soybean varieties qualifies as biomarkers based on their phenotypic traits.

Analysis of the data collected was done using four software: SIFT, Polyphen, PANTHER, and I-mutant 2.0, which are designed to detect the rate of functional and structural variations caused by the nsSNPs in cultivated and wild soybean varieties. Genotypic information obtained in the analysis was used to develop a core collection of biomarkers based on whether nsSNP content was found in more than half of the 16 samples. Therefore, the list of biomarker candidates developed from this study showed that Korean soybean could provide valuable information needed in both future crop genetic research and identification of biomarkers.

## INTRODUCTION

Soybean is a self-pollinating species with severe genetic bottlenecks during domestication. According to Hyten et al. (2006), approximately 81% of the rare alleles are lost during domestication. Soybeans have relatively low nucleotide polymorphism rates than other crop species (Haun et al., 2011). There is a wide range of phenotypic variation between wild and cultivated soybeans (Fasoula et al., 2005). These genetic variations are associated with a change in a single nucleotide found in the DNA.

A change in single nucleotide polymorphism (SNP) in the DNA of soybean leads to genetic variation between individuals within the soybean species. Single nucleotide polymorphism (SNP) in soybean does not occur frequently. However, SNP with a wide range of heritable phenotypic differences is neutral (Haun et al., 2011). Non-synonymous single nucleotide polymorphisms (nsSNPs) occurring in coding or regulatory regions of the DNA alter both protein structure and protein function. For example, wild soybean (G. soja) is different from cultivated soybean (G. max) in terms of flowering time and grain weight.

Both wild and domestic soybeans have 20 chromosomes, almost the same amino acid sequences, but a varying number of nsSNPs. These phenotype alterations may result from nsSNPs which affect gene expression and alter splice sites, leading to new gene products. Structural variations between the two genomes include deletion, insertions, inversions, and translocation. These nsSNPs introduce or expel unexpected charged residue or inflexible amino acids with benzene ring (s) or some amino acids, which significantly alter the residue (Bromberg et al., 2007). The deleterious effects of nsSNPs in soybeans, therefore, make them probable biomarker candidates.

This argument prompted the extraction of a sizeable nsSNP set from 16 soybean varieties, both wild and cultivated. The collection comprised of 6 wild varieties which include (IT162825, IT178480, IT182869, IT182840, IT182848, and IT182932) and 10 cultivated varieties (Sowon, Peuren, Kangyo, Ilpumgeomjeong, Seoritae, PI96983, Haman, Kumjeongol, Hwangkeum, and Williams 82k)]. Cultivar William 82 was used as the genetic sequence reference unit to compare it with the sequence of the collection of 16 cultivated soybean varieties in Korea. According to Haun et al., (2011), the genetic heterogeneity in Williams 82 soybean primarily originated from the differential segregation of polymorphic chromosomal regions. According to Haun et al., (2011), this differential segregation occurred during the hybridization of the backcross and single-seed descent generations over the years.

Although there are many wet-lab experiments done on the functional impact of nsSNPs, it has proven difficult to investigate all of them since they are believed to be more than 100000. However, bioinformatics approaches have increased the prediction of the functional effect of nsSNPs. This study predicts phenotypic consequences of amino acid substitution such as seed composition, seed weight, seed size, and plant height using computational methods in silico software such as SIFT (Sorting Intolerant from Tolerant) (Ng et al., 2003), Polyphen (Polymorphism Phenotyping) (Ramenskyet al., 2002), PANTHER (Protein Analysis Through Evolutionary Relationships) (Thomas et al., 2003), and I-Mutants.

This methodology has been proven to develop a more prospective list of biomarker candidates, which will be needed for future QTL investigations (Di et al., 2009). The limitation of using computational methods is that they do not always guarantee the accuracy of real biomarkers due to the severe dependence on their exclusively theoretical judgments. However, it facilitates the selection of a core list of biomarker candidates from a pool of a large population of available data (Bromberg et al., 2007). Therefore, the primary purpose of this paper is to present a more specific list of nsSNPs, which are responsible for the structural and functional alteration of the genetic material of soybeans during domestication. The study employed the use of 16 varieties selected from wild and cultivated soybeans through gene sequencing to derive a list of biomarkers based on the rate of nsSNPs impact on their gene structure.

## MATERIALS AND METHODS

A total of 16 soybean varieties and their nsSNPs composition data obtained from the Korean Bioinformation Center (KOBIC) was used for this study. Genotype analysis was done to isolate soybean varieties with nsSNPs as probable biomarker candidates. Four computing software were used to examine the nsSNP data and determine whether they qualify as biomarkers. This software included SIFT, PolyPhen, PANTHER, and I-Mutants. Genetic diversity analysis was performed in the LINUX environment at the bio-evaluation center in the Korea Research Institute of Bioscience and Biotechnology. A summary of the software functions is as follows.

1. A SIFT software (https://sift.bii.a-star.edu.sg): It is known to distinguish between functionally tolerated and deleterious amino acid changes in mutagenesis studies. It predicts whether amino acid substitution affects protein function or potentially alters the phenotype of genes occurring in the varieties. This is based on sequence homology (the degree of conservation of amino acid residues in sequence alignments derived from closely related courses, collected through PSI-BLAST) and the physical properties of amino acids, resulting in either “Tolerated (more than 0.05) “or “Deleterious (less than 0.05)”. To be brief, SIFT presumes that essential amino acids will be conserved in the protein family, and so changes at well-conserved positions tend to be predicted as “deleterious.”
2. A Polyphen-2 software (http://genetics.bwh.harvard.edu/pph2): It is a new development of the Polyphen tool for annotating coding nsSNPs. It determines the properties of nsSNPs based on a combination of phylogenetic, structural, and sequence annotation information characterizing a substitution and its position in the protein. The Polyphen values range from 0 (tolerated) and 1 (most likely to be deleterious) according to the degree of structural change. Mapping of amino acid replacement to the known 3D structure reveals whether the alternative is possible to destroy the hydrophobic core of a protein, electrostatic interactions, interactions with ligands, or other essential protein features. If a query protein’s spatial structure is unknown, one can use the homologous proteins with known structure (Adzhubei et al., 2010).
3. A PANTHER software (http://www.pantherdb.org): It predicts the functional impact of the related nsSNPs in the absence of direct experimental evidence based on published scientific empirical evidence and evolutionary relationships (Thomas et al., 2003). The PANTHER values range from 0 (neutral) to (approx.) −10 (most likely to be deleterious) according to the degree of functional impairment. The PSCI (substitution position-specific evolutionary conservation) values where less than −3 and the deleterious probability of more than 0.5 are assumed to be “deleterious” [-3 of PSCI equals to 0.5 of probability deleterious].
4. An I-Mutant 2.0 software (http://folding.biofold.org/i-mutant/i-mutant2.0.html): It checks the stability of the proteins based on the analysis for solvent accessibility (NetASA) and secondary structure (DSSP). Nature generally prefers stable conditions. Values less than zero are assumed to be “unstable” or “deleterious.” So if the nsSNPs involved make any protein structure unstable, they will be classified as “deleterious” and are presumed to have individual aims to exist.

## RESULTS

The study considered multiple ways of nsSNP impacts on protein structure and functions in 16 soybean varieties drawn from wild and domestic varieties found in Korea. SNPs with the highest likelihood of being functionally relevant were the most suitable samples to be examined and ranked based on SIFT, PolyPhen, PANTHER, and I Mutant 2.0 scores. The study selected 91 most likely nsSNP candidates that had an impact on the protein structures and functions of the soybeans (see APPENDIX).

Table 1 shows the properties of nsSNPs simulated by SIFT and Polyphen for the three cases of nsNSPs exclusive in wild varieties (B), nsSNPs unique in cultivated varieties (A), and nsSNPs simultaneously occurring in both wild and cultivated varieties (AB). In (B), out of 39258 nsSNPs (13003 genes), 5175 nsSNPs were observed to have a potentially damaging effect on the protein function of the genome, whereas 901 nsSNPs were finally chosen through empirical intuition that nsSNPs appearing in more than half of the wild varieties are functionally important. In (A), out of 4302 nsSNPs (2118 genes), 1146 nsSNPs were observed to have a potentially damaging effect, whereas 77 nsSNPs were finally chosen with the same reason. Similarly, in (AB), out of 131992 nsSNPs (20595 genes), 20361 nsSNPs were observed to have potentially damaging effects, and 2318 nsSNPs were added to the core collection list that would be significant for the study.

**Table 1.** The properties of nsSNPs simulated by SIFT and Polyphen.

|  | **SIFT**<br>**(B)** | **Polyphen**<br>**(B)** | **SIFT**<br>**(A)** | **Polyphen**<br>**(A)** | **SIFT**<br>**(AB)** | **Polyphen**<br>**(AB)** |
| --- | --- | --- | --- | --- | --- | --- |
| **Gene** | 13003 |  | 2118 |  | 20595 |  |
| **nsSNP** | 39258 (100%) |  | 4302 (100%) |  | 131992 (100%) |  |
| **Deleterious** | 9655<br>(25%) | 10952<br>(28%) | 1697<br>(39%) | 1709<br>(40%) | 40641<br>(31%) | 32582<br>(25%) |
| **Deleterious**<br>**in both SIFT**<br>**and**<br>**Polyphen** | 1≤nsSNP≤6 |  | 1≤nsSNP≤10 |  | 1≤nsSNP≤16 |  |
|  | 5175 (13%) |  | 1146 (27%) |  | 20361 (15%) |  |
|  | 3≤nsSNP≤6 |  | 5≤nsSNP≤10 |  | 8≤nsSNP≤16 |  |
|  | 901 (2%) |  | 77 (2%) |  | 2318 (2%) |  |
| **Tolerated** | 22117<br>(58%) | 18488<br>(47%) | 2340<br>(54%) | 1780<br>(41%) | 79653<br>(60%) | 53611<br>(41%) |
| **Unknown** | 7486 | 9818 | 265 | 813 | 5450 | 45799 |
| # B: a set of genes with nsSNPs exclusive in wild varieties, A: a set of genes with nsSNPs unique in cultivated varieties, and AB: a set of genes with nsSNPs occurring simultaneously in both wild and cultivated varieties.<br># The maximum number of nsSNPs per variation position of each gene:<br>B (6), A(10), and AB (16) |  |  |  |  |  |  |

The selection of nsSNPs from these three different cases were further analyzed using software PANTHER (more than 0.98) and I-mutant (less than 0 when pH=7 and T= 298K), in an aim to develop a more accurate shorter list of most likely and tractable candidates of nsSNPs. The study further examined the gene flow strength of these nsSNP data. F_ST_ [Wright’s fixation index (1951), an estimator of the amount of gene flow between varieties] value of each gene was added to the nsSNP data. The F_ST_ was calculated as follows:

F_ST_ in a gene = The number of nsSNPs generated by the comparison of two genes selected from two subsets [one gene in 6 wild soybean subset and the other in 11 cultivated soybean subset (including W 82)] / The number of nsSNPs generated by the comparison of two genes selected from a set of 17 wild and cultivated soybeans.

Based on a computational study of these nsSNP data, agronomically significant 91 biomarker candidates were shortlisted from the three cases of (B), (A), and (AB). In the subsequent studies, 3-D structural analysis and molecular dynamics simulation could be applied in assessing the functional properties and specificity of 91 biomarkers to check whether they are realistic.

## CONCLUSION

The study observed that it is possible to identify the impact of nsSNPs on protein structure and function of genes in mapping biomarkers using PolyPhen, SIFT, I-mutant 2.0 PANTHER computer software. The application of bioinformatic software and the interpretation of sample soybean data were subjectively adopted, and most of them may not give a realistic picture. However, some of them have a higher probability of being recognized as agronomically essential biomarkers in future. Further information on nsSNPs can be obtained from public databases (i.e., http://www.soybase.org and http://www.ncbi.nlm.nih.gov) to find out whether some nsSNPs in the list were already reported as biomarkers or not.

The biomarker list developed using nsSNP levels in such wild and cultivated soybean varieties will facilitate future research in the following ways:

1. To establish essential methodologies to check structural changes during hybridization
2. In the extraction of biologically relevant nsSNPs with phenotypic traits favourable to enhance agronomical characteristics and improve the social economy through increased production.
3. To facilitate tracking of the domestication process and gene manipulation in the population genetics.

Data on gene sequencing has been observed to change with time continuously. The information on nsSNPs provided in our list of biomarkers may need to be revised in future to ensure relevance. For further reference, all the biomarker lists and source codes will be provided upon request.

## ACKNOWLEDGEMENTS

The authors are grateful to the bio-evaluation center at the Korea Research Institute of Bioscience and Biotechnology for providing us with the nsSNP data of 16 Korean soybean varieties used in this study.

## APPENDIX A shortlist of biomarker candidates

Note that the number in () of substitution denotes the frequency of the substitution at the specified nucleotide, whereas the values of PANTHER more than 0.98 indicate “extremely deleterious”.

**(1) biomarker candidates assigned to A (Ch = Chromosome)**

(1) 1 biomarker candidates assigned to A (Ch = Chromosome)
| Ch. No | Gene Name | Substitution | SIFT | Polyphen | PANTHER | I-mutant 2.0 | SNP-F<sub>ST</sub> |
| --- | --- | --- | --- | --- | --- | --- | --- |
| 15 | Glyma15g17100.1 | P19T(7) | 0 | 1 | 0.98 | -1.29 | 0.4 |

**(2) 15 biomarker candidates assigned to B**

| Ch. No | Gene Name | Substitution | SIFT | Polyphen | PANTHER | I-mutant2.0 | SNP-F<sub>ST</sub> |
| --- | --- | --- | --- | --- | --- | --- | --- |
| 11 | Glyma11g33430.1 | C29R(5) | 0 | 1 | 1 | -0.24 | 0.09 |
| 10 | Glyma10g33120.1 | L272H(5) | 0 | 1 | 1 | -3.04 | 0.11 |
| 8 | Glyma08g01870.1 | K202T(3) | 0 | 1 | 1 | -1.43 | 0.14 |
| 8 | Glyma08g01870.2 | K202T(3) | 0 | 1 | 1 | -1.43 | 0.14 |
| 8 | Glyma08g01870.3 | K202T(3) | 0 | 1 | 1 | -1.43 | 0.14 |
| 17 | Glyma17g13910.1 | Q136N(3) | 0 | 1 | 1 | -1.42 | 0.22 |
| 11 | Glyma11g33430.1 | P77L(5) | 0.01 | 1 | 0.99 | -1.19 | 0.09 |
| 2 | Glyma02g41690.1 | F221M(3) | 0 | 1 | 0.99 | -0.02 | 0.15 |
| 6 | Glyma06g05670.1 | S84C(3) | 0.01 | 1 | 0.99 | -2.04 | 0.21 |
| 17 | Glyma17g04720.1 | E125G(3) | 0 | 1 | 0.99 | -0.77 | 0.21 |
| 8 | Glyma08g00520.1 | I78H(6) | 0 | 1 | 0.98 | -2.56 | 0 |
| 3 | Glyma03g26890.1 | S365A(3) | 0 | 1 | 0.98 | -0.66 | 0.08 |
| 11 | Glyma11g33430.1 | S96Y(5) | 0.01 | 0.99 | 0.98 | -1.16 | 0.09 |
| 3 | Glyma03g31400.1 | F152P(4) | 0 | 1 | 0.98 | -2.09 | 0.15 |
| 13 | Glyma13g21980.1 | G153D(4) | 0.01 | 1 | 0.98 | -2.43 | 0.15 |

**(3) 75 biomarker candidates assigned to AB**

| Ch. No | Gene Name | Substitution | SIFT | Polyphen | PANTHER | I-mutant2.0 | SNP-F<sub>ST</sub> |
| --- | --- | --- | --- | --- | --- | --- | --- |
| 3 | Glyma03g12230.1 | R396W(10) | 0 | 1 | 1 | -1.08 | 0.36 |
| 6 | Glyma06g45710.1 | R466W(8) | 0 | 1 | 1 | -0.78 | 0.45 |
| 8 | Glyma08g01890.1 | R36A(9) | 0 | 1 | 1 | -0.53 | 0.54 |
| 8 | Glyma08g01890.2 | R36A(9) | 0 | 1 | 1 | -0.53 | 0.54 |
| 8 | Glyma08g28170.1 | I347P(12) | 0 | 1 | 1 | -2.57 | 0.54 |
| 14 | Glyma14g05280.1 | L6S(14) | 0 | 1 | 1 | -1.52 | 0.54 |
| 3 | Glyma03g07910.1 | F233A(16) | 0 | 1 | 1 | -2.83 | 0.57 |
| 1 | Glyma01g37330.1 | L372T(16) | 0 | 1 | 1 | -3.19 | 0.62 |
| 1 | Glyma01g41090.1 | D77T(16) | 0 | 1 | 1 | -1.7 | 0.63 |
| 1 | Glyma01g41090.1 | K167G(16) | 0 | 1 | 1 | -1.35 | 0.63 |
| 1 | Glyma01g41090.1 | G80D(16) | 0 | 1 | 1 | -1.12 | 0.63 |
| 1 | Glyma01g41090.1 | T78Q(16) | 0 | 1 | 1 | -0.87 | 0.63 |
| 1 | Glyma01g41090.1 | S165T(16) | 0 | 1 | 1 | -0.15 | 0.63 |
| 1 | Glyma01g41090.1 | A166L(16) | 0 | 1 | 1 | -0.15 | 0.63 |
| 10 | Glyma10g42260.1 | F306E(8) | 0 | 1 | 0.99 | -1.85 | 0.24 |
| 2 | Glyma02g03450.1 | G109S(9) | 0 | 1 | 0.99 | -1.46 | 0.26 |
| 18 | Glyma18g07340.1 | S343C(8) | 0.03 | 0.69 | 0.99 | -1.49 | 0.38 |
| 8 | Glyma08g13040.1 | K1196N(10) | 0 | 1 | 0.99 | -0.65 | 0.38 |
| 16 | Glyma16g17430.1 | N263G(10) | 0.01 | 1 | 0.99 | -1.6 | 0.41 |
| 8 | Glyma08g13040.2 | K52N(10) | 0 | 1 | 0.99 | -0.65 | 0.47 |
| 13 | Glyma13g15350.1 | G1772R(11) | 0.04 | 0.97 | 0.99 | -0.37 | 0.49 |
| 18 | Glyma18g46900.1 | R102W(11) | 0 | 1 | 0.99 | -0.68 | 0.49 |
| 14 | Glyma14g11610.1 | G320V(8) | 0.02 | 0.9 | 0.99 | -0.27 | 0.51 |
| 20 | Glyma20g35610.1 | V292E(10) | 0 | 0.99 | 0.99 | -0.55 | 0.52 |
| 18 | Glyma18g50290.1 | S108C(9) | 0.04 | 1 | 0.99 | -0.81 | 0.52 |
| 13 | Glyma13g25920.1 | I485H(14) | 0.01 | 1 | 0.99 | -3.14 | 0.52 |
| 20 | Glyma20g23360.1 | I120Y(10) | 0 | 0.99 | 0.99 | -2.28 | 0.53 |
| 9 | Glyma09g41840.1 | L285C(10) | 0 | 1 | 0.99 | -2.65 | 0.56 |
| 4 | Glyma04g09630.1 | W520G(16) | 0 | 1 | 0.99 | -1.72 | 0.56 |
| 2 | Glyma02g03200.1 | W153G(15) | 0 | 1 | 0.99 | -4.15 | 0.57 |
| 2 | Glyma02g03200.1 | F161S(15) | 0 | 1 | 0.99 | -3.03 | 0.57 |
| 7 | Glyma07g04560.1 | G64R(14) | 0 | 1 | 0.99 | -1.6 | 0.57 |
| 9 | Glyma09g08900.1 | L421P(13) | 0 | 1 | 0.99 | -0.08 | 0.6 |
| 2 | Glyma02g26410.1 | L345D(16) | 0 | 1 | 0.99 | -2.24 | 0.6 |
| 12 | Glyma12g35580.1 | P653G(16) | 0 | 1 | 0.99 | -1.66 | 0.61 |
| 12 | Glyma12g35580.1 | L650R(16) | 0 | 1 | 0.99 | -1.3 | 0.61 |
| 12 | Glyma12g35580.1 | M625R(16) | 0 | 0.98 | 0.99 | -0.68 | 0.61 |
| 9 | Glyma09g08910.1 | V318R(15) | 0 | 1 | 0.99 | -2.39 | 0.61 |
| 1 | Glyma01g41090.1 | G169H(16) | 0 | 1 | 0.99 | -2.73 | 0.63 |
| 1 | Glyma01g41090.1 | I179Q(16) | 0 | 1 | 0.99 | -1.39 | 0.63 |
| 14 | Glyma14g21230.2 | G18E(16) | 0 | 1 | 0.99 | -0.28 | 0.63 |
| 14 | Glyma14g21230.3 | G18E(16) | 0 | 1 | 0.99 | -0.28 | 0.63 |
| 17 | Glyma17g20110.1 | F448V(10) | 0 | 1 | 0.99 | -1.77 | 0.88 |
| 15 | Glyma15g12430.1 | V133H(10) | 0 | 1 | 0.98 | -3.17 | 0.33 |
| 17 | Glyma17g09310.1 | P91T(9) | 0 | 1 | 0.98 | -0.97 | 0.33 |
| 12 | Glyma12g22640.1 | F158S(9) | 0 | 0.97 | 0.98 | -3.43 | 0.46 |
| 17 | Glyma17g05650.1 | S645D(12) | 0 | 1 | 0.98 | -0.31 | 0.5 |
| 18 | Glyma18g45530.1 | V335R(8) | 0 | 1 | 0.98 | -1.78 | 0.51 |
| 3 | Glyma03g03460.1 | L472Q(12) | 0 | 1 | 0.98 | -1.47 | 0.52 |
| 3 | Glyma03g03460.1 | G176L(12) | 0.03 | 1 | 0.98 | -0.58 | 0.52 |
| 13 | Glyma13g15590.1 | W129M(10) | 0 | 1 | 0.98 | -1.04 | 0.52 |
| 1 | Glyma01g43610.1 | F481K(10) | 0 | 1 | 0.98 | -1.91 | 0.53 |
| 1 | Glyma01g43610.1 | R115C(10) | 0 | 1 | 0.98 | -0.94 | 0.53 |
| 3 | Glyma03g34130.1 | R107W(10) | 0 | 1 | 0.98 | -0.74 | 0.53 |
| 1 | Glyma01g42120.1 | L33S(12) | 0 | 0.86 | 0.98 | -2.08 | 0.55 |
| 9 | Glyma09g41840.1 | G296H(11) | 0 | 1 | 0.98 | -2.38 | 0.56 |
| 9 | Glyma09g41840.1 | V364W(10) | 0 | 1 | 0.98 | -1.72 | 0.56 |
| 9 | Glyma09g41840.1 | E366K(10) | 0 | 1 | 0.98 | -0.97 | 0.56 |
| 9 | Glyma09g41840.1 | V298M(10) | 0 | 1 | 0.98 | -0.3 | 0.56 |
| 4 | Glyma04g09630.1 | W521G(16) | 0 | 1 | 0.98 | -2.08 | 0.56 |
| 2 | Glyma02g03200.1 | T157R(15) | 0.01 | 1 | 0.98 | -0.28 | 0.57 |
| 13 | Glyma13g29580.1 | D143T(14) | 0.04 | 0.75 | 0.98 | -0.55 | 0.57 |
| 13 | Glyma13g29580.1 | A268R(14) | 0 | 0.87 | 0.98 | -0.5 | 0.57 |
| 3 | Glyma03g07910.1 | Y230E(16) | 0 | 0.99 | 0.98 | -0.43 | 0.57 |
| 9 | Glyma09g08900.1 | Y428R(13) | 0 | 1 | 0.98 | -0.26 | 0.6 |
| 12 | Glyma12g35580.1 | A623Q(16) | 0 | 1 | 0.98 | -0.86 | 0.61 |
| 9 | Glyma09g08910.1 | V320G(15) | 0 | 1 | 0.98 | -3.71 | 0.61 |
| 9 | Glyma09g08910.1 | G331D(15) | 0 | 1 | 0.98 | -0.73 | 0.61 |
| 15 | Glyma15g13620.1 | L303G(16) | 0 | 1 | 0.98 | -4.14 | 0.63 |
| 6 | Glyma06g10420.1 | W246P(16) | 0 | 1 | 0.98 | -2.15 | 0.63 |
| 15 | Glyma15g13620.1 | H307Q(16) | 0 | 0.99 | 0.98 | -1.5 | 0.63 |
| 6 | Glyma06g10420.1 | Y235R(16) | 0 | 1 | 0.98 | -0.88 | 0.63 |
| 1 | Glyma01g41090.1 | Q81R(16) | 0 | 1 | 0.98 | -0.44 | 0.63 |
| 1 | Glyma01g41090.1 | C219S(16) | 0 | 1 | 0.98 | -0.04 | 0.63 |
| 1 | Glyma01g41090.1 | E163Q(16) | 0 | 1 | 0.98 | -0.03 | 0.63 |

## REFERENCES

David L. Hyten, Qijian Song, Youlin Zhu, Ik-Young Choi, Randall L. Nelson, Jose M. Costa, James E. Specht, Randy C. Shoemaker, Perry B. Cregan. 2006. Impacts of genetic bottlenecks on soybean genome diversity. PANS 103: 16666–16671.

William J. Haun, David L. Hyten, Wayne W. Xu, Daniel J. Gerhardt, Thomas J. Albert, Todd Richmond, Jeffrey A. Jeddeloh, Gaofeng Jia, Nathan M. Springer, Carroll P. Vance, Robert M. Stupar. 2011. The Composition and Origins of Genomic Variation among Individuals of the Soybean Reference Cultivar Williams 82. Plant Physiology Preview 155: 645–655.

Fasoula VA and Boerma HR. 2005. The divergent selection at ultra-low plant density for seed protein and oil content within soybean cultivars. Field Crops Res. 91: 217–229.

Yana Bromberg and Burkhard Rost. 2007. SNAP: predict the effect of non-synonymous polymorphisms on function. Nucleic Acids Research 35: 3823–3835.

Pauline C. Ng and Steven Henikoff. 2003. SIFT: predicting amino acid changes that affect protein function. Nucleic Acids Research 31: 3812–3814.

Ramensky V, Bork P, Sunyaev S. 2002. Human non-synonymous SNPs: server and survey. Nucleic Acids Res 30:3894–3900

Paul D. Thomas, Michael J. Campbell, Anish Kejariwal, Huaiyu Mi, Brian Karlak, Robin Daverman, Karen Diemer, Anushya Muruganujan, Apurva Narechania. 2003. PANTHER: A Library of Protein Families and Subfamilies Indexed by Function. Genome Res. 13: 2129–2141.

Emidio Capriotti, Piero Fariselli, Rita Casadio. 2005. I-Mutant2.0: predicting stability changes upon mutation from the protein sequence or structure. Nucleic Acids Research. 33: W306–W310.

Yuan Ming Di, Eli Chan, Ming Qian Wei, Jun-Ping Liu, Shu-Feng Zhou. 2009. Prediction of Deleterious Non-synonymous Single-Nucleotide Polymorphisms of Human Uridine Diphosphate Glucuronosyltransferase Genes. The AAPS Journal. 11: 460–480.

Adzhubei IA, Schmidt S, Peshkin L, Ramensky VE, Gerasimova A, Bork P, Kondrashov AS, Sunyaev SR. 2010. A method and server for predicting damaging missense mutations. Nat Methods 7: 248–249.

Wright S (1951). The genetical structure of populations. Ann Eugen 15: 323–354.

